# Fabrication of size-coded amphiphilic particles with a configurable 3D-printed microfluidic device for the formation of particle-templated droplets

**DOI:** 10.1101/2023.09.20.558669

**Authors:** Muhammad Usman Akhtar, Mehmet Akif Sahin, Helen Werner, Ghulam Destgeer

**Affiliations:** Control and Manipulation of Microscale Living Objects, TranslaTUM - Center for Translational Cancer Research, Department of Electrical Engineering, School of Computation, Information and Technology, Technical University of Munich, Germany

**Keywords:** 3D printing, microfluidics, stop-flow lithography, amphiphilic particles

## Abstract

Compartmentalizing an aqueous media into numerous nanoliter-scale droplets has substantially improved the performance of amplification assays. Particle-templated droplets or *dropicles* offer a user-friendly workflow for creating uniform volume compartments upon simple mixing of reagents and particles by using common laboratory apparatus. Amphiphilic shape-coded particles have been demonstrated to spontaneously hold aqueous droplets within hydrophilic cavities for multiplexed diagnostic assays. Here, we have proposed a configurable 3D-printed microfluidic device for the tunable fabrication of amphiphilic size-coded particles. The device was configured with multiple outlet tubings of different diameters and photomasks of variable slit lengths to fabricate a wide range of size-coded particles. We have fabricated >10 unique particle codes using a single reconfigurable device. The cross-sectional profile of the particles was further engineered by tuning the flow rate ratios of precursor streams to vary the inner and outer diameters of the particles and the thicknesses of the inner hydrophilic and outer hydrophobic layers. A range of cavity diameters and particle lengths enabled *dropicle* volumes of ∼1nL to ∼30nL. The fabricated particles were characterized by their ability to hold uniform droplet volumes and to orient themselves facing upwards or sideways in a well plate based on their aspect ratios.

## 1. Introduction

Biochemical samples are compartmentalized within small containers to spatially distribute the sample volume and perform numerous parallel reactions with minimal crosstalk. ^1,2^ The ability to form and analyze a large number of uniform reaction volumes is critical for several diagnostic assays,^3,4^ such as quantitative polymerase chain reaction (PCR), single molecule detection in digital bioassays,^5^ swarm sensing in analog assays,^6^ etc. The compartments uniformly distribute and confine analytes of interest for signal accumulation, amplification, and quantification.^7^ A solid anchoring surface within these compartments provides analyte-specific binding sites, which are required for capturing the target molecules and performing intermittent washing steps in an assay. Moreover, the compartments can be labeled individually for multiplexing to identify multiple analytes simultaneously.^8^ Conventionally, test tubes and micro-well plates have been commonly used as compartmentalized biochemical reactors, but they are limited by the microliter-scale reaction volumes and the number of parallel compartments. ^1,2^

Microfluidic droplet generators have been extensively employed in the past two decades to produce uniform sub-nanoliter reaction volumes with the ability to encapsulate single biomolecules or cells. ^9-11^ The absence of a solid anchoring surface within the aqueous droplets limits their applications; therefore, colored microbeads are simultaneously encapsulated with the analytes to introduce binding sites and multiplexing capabilities.^12^ Microfluidics have led to advances in digital PCR, enzyme-linked immunosorbent assay (ELISA), and single-cell secretion assays. ^13^ Despite having demonstrated great potential, microfluidic droplet-based compartmentalization faces challenges in its widespread adoption, mainly because of the need for advanced cleanroom facilities for device fabrication and specialized instruments to operate these devices by experienced personnel.

The particle-templated droplets or *dropicles* have emerged as a simplified, user-friendly, and instrument-free workflow for uniform volume compartmentalization. ^14^ Typically, the *dropicles* are formed by simply mixing the pre-cured solid particles with the aqueous and oil media inside a centrifuge tube or a well plate by pipetting. An aqueous droplet is templated by the solid particle surrounded by the continuous oil phase. Single ^15^ and multi-material particles ^16-18^ with different shapes, such as spherical,^14^ crescent,^19^ horseshoe,^20^ hollow cylindrical,^16-18^ etc., have been fabricated by using flow lithography techniques for monodisperse droplets formation.^21^ Particularly, the co-axial flow lithography method, utilizing 3D printed microfluidic devices, has shown great flexibility in spatially modulating the cross-sectional flow profiles to fabricate shape-coded hollow particles for multiplex diagnostics applications.^16^

In this study, we have reported a configurable 3D-printed microfluidic device that can be used with multiple outlet tubings and photomasks to fabricate cylindrical particles with a range of inner and outer diameters, lengths and cavity volumes. The microfluidic device and the photomasks were 3D printed using an in-house rapid prototyping protocol to significantly reduce the costs associated with outsourcing. For the particle fabrication, we have demonstrated a stop-flow lithography ^22,23^ setup utilizing a rapidly switchable LED-based light source to instantaneously control the UV exposure of curable polymer precursors without the need for a mechanical shutter. ^24^ The 3D-printed microfluidic device effectively sculpted the precursor flow streams in an annular cross-sectional profile, while the selective exposure of UV light controlled the length of particles, thereby providing an on-demand control over the radial and axial dimensions of the fabricated particles. Moreover, by controlling the flow rate ratios of individual streams, we modulated the thickness of the hydrophilic and hydrophobic layers, and inner and outer diameters of the amphiphilic particle. All the fabricated size-coded particles were characterized for their ability to hold uniform aqueous droplets. The aspect ratio of the particles was critical in determining orientation of the particle and uniformity of templated droplet. We carefully adjusted the cavity diameter and length to realize a constant droplet volume for different size-coded particles. Our stop-flow lithography setup with the configurable device enabled a modular batch-based fabrication process that utilized the maximum flow length and exposure area for high throughput production of particles.

The configurable 3D microfluidic device had 4-coaxially stacked microchannels, each connected with a separate inlet for the injection of polymer precursor into the device (**Figure 1**A). The polymer precursors, comprising inert hydrophobic, curable hydrophobic, curable hydrophilic, and inert hydrophilic streams, entered the device at a flow rate of *Q*_l_, *Q*_2_, *Q*_3_, and *Q*_4_, respectively. The channels had concentric cross-sectional profiles with a tapered nozzle at the end to sequentially squeeze the multilayered stacked streams into a transparent tubing connected at the outlet port. The interior surface of the microfluidic device at the far-right end was a threaded fastener to connect the device with the transparent tubing through a nut and ferrule of a flangeless fitting for easy replacement. The concentric streams in the transparent tubing were shined by UV light through a 3D printed photomask (Figure 1B). By connecting outlet tubings with different inner diameters to the same 3D-printed device, diameter coded (*D*-coded) amphiphilic particles were fabricated with distinct outer diameters (*D*_o_) (Figure 1C). By varying the slit length of the photomasks, we fabricated length coded (*L*-coded) particles with different lengths (*L*). The flow rate ratios of 4-streams were modulated to control the cavity diameter (*D*_*c*_) and the thicknesses of inner (*t*_*in*_) and outer (*t*_*out*_) layers of the particles (Figure 1D). Droplets were encapsulated in the cavity of the amphiphilic particles after two medium exchange steps: (1) from ethanol to phosphate buffer saline (PBS), and (2) from PBS to oil (Figure 1E). A fluorescent dye was mixed in the PBS solution for the visualization of the encapsulated droplets. An aqueous droplet was captured in the hydrophilic cavity of each particle as the added oil interacted with the outer hydrophobic layer of the particle (Figure 1F).

**Figure 1.**
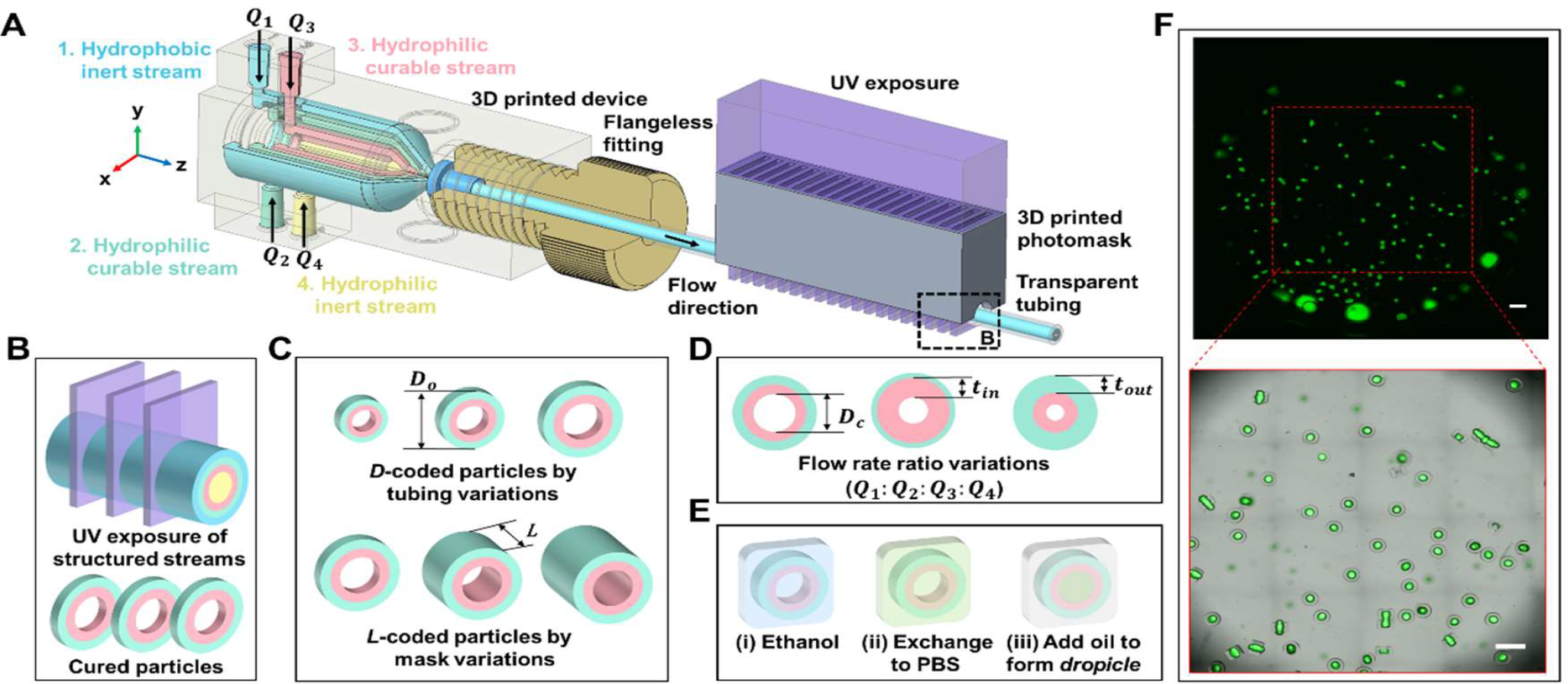
Fabrication of concentric size-coded amphiphilic particles through a configurable 3D printed device. (A) Schematic of the configurable 3D printed coaxial flow microfluidic device with a flangeless fitting connector at the outlet. A 3D printed photomask positioned on top of the transparent tubing for selective UV exposure. (B) Enlarged view of the UV exposure of the four concentric streams in the tubing through the photomask for curing amphiphilic particles with hydrophobic outer and hydrophilic inner layers. (C) Size-coded amphiphilic particles included diameter- and length-based *D*-coded and *L*-coded particles, respectively. (D) The flow rate ratio variations for tuning the cavity diameter (*D*_*c*_) and the thickness of inner (*t*_*in*_) and outer (*t*_*out*_) layers of particle. (E) Dropicle formation protocol: (i) particles in ethanol seeded in the well plate, (ii) medium exchanged from ethanol to PBS, (iii) add oil to encapsulate *dropicle*. (F) A full-well image of a 24-well plate with several *dropicles* depicted in the fluorescent (top) and fluorescent overlaid brightfield (bottom) channels (Scale bar 1mm).

## 2. Results and Discussion

### 2.1 Design and 3D printing of the microfluidic device

A concentric 3D microfluidic device with a coaxial geometry of four channels was designed with a tapered nozzle and a threaded connection (**Figure 2A**). The innermost channel of the device had a diameter of 0.8mm, while the outer annular channels had a wall-wall gap of 0.6mm, with a wall thickness of 0.5mm in between two stacked channels. Each channel sequentially squeezed the fluid through a nozzle toward the outlet port of the device with a diameter of 0.7mm. The device was configurable with different outlet tubings having the same outer diameter (*OD*_*t*_) of 1.6mm and different inner diameters (*ID*_*t*_) of 0.5mm, 0.75mm, and 1.0mm. The tubing was connected to the device through the ferrule and nut of a flangeless fitting. We performed ‘computation fluid dynamics’ simulations to confirm that the designed device could effectively sculpt the polymer streams in a desired concentric cross-sectional flow profile (**Figure S1**A). Thereby, the fabrication of the designed device was initiated. We used a resin-based 3D printer (vat polymerization technology) for rapid prototyping of our design due to their higher resolution and lower cost. 3D printing of open microfluidic channels with dimensions <100μm has been demonstrated with high-resolution resin-based 3D printers.^25^ However, 3D printing our fully enclosed microfluidic device proved tricky because of undesired curing of resin filled inside the hollow channels due to repeated exposure to leaky UV light. To avoid clogging the 3D printed channels, we have adopted the device design to be compatible with the commercially available low-cost resins for 3D printing. We enlarged the channel gaps to more than 400μm and adjusted the angle of tapered nozzles for easy drainage of uncured resin from the channels. Moreover, we used the minimal UV exposure settings suitable for the resin to avoid undesired overexposure of the resin. Our fully enclosed microfluidic device with several intertwined microchannels was successfully 3D printed at a very low cost of around $1.3 (Figure 2B and **Table S1**). The dimensions of the designed and 3D-printed device matched very well, whereas a slight reduction in all the dimensions was attribute to the shrinkage of the cured resin after post curing (see supporting information for details).

**Figure 2.**
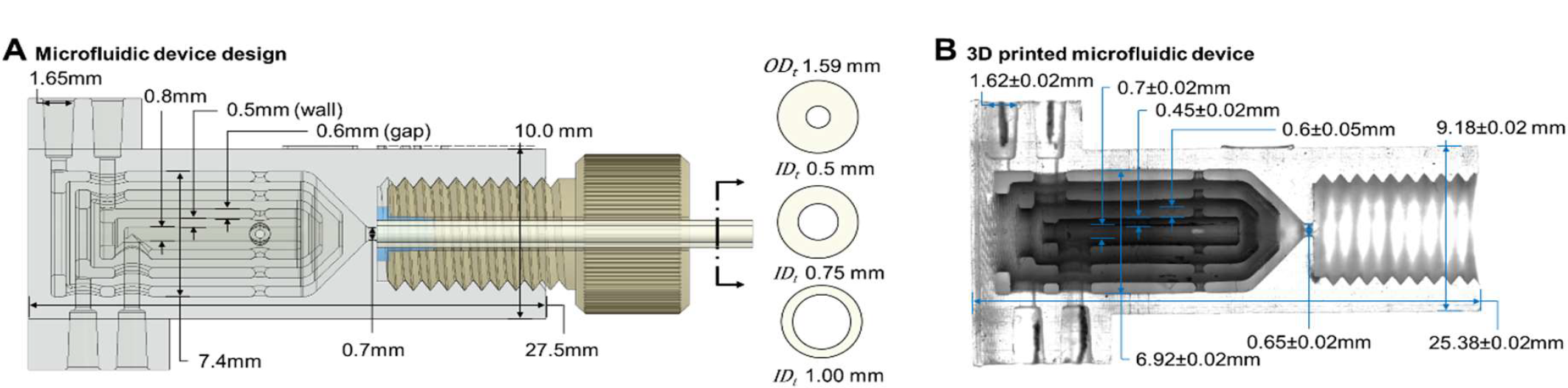
Configurable microfluidic device. (A) Design of a microfluidic device with four inlets, each connected to a channel separated by a wall. The channels merged at the threaded outlet for a screw-connection of transparent tubing with the flangeless fitting. Three different tubings were used here, each with an outer diameter (*OD*_*t*_) of 1.59mm and an inner diameter (*ID*_*t*_) of 500μm, 750μm, and 1000μm. (B) Half cross-section of the 3D printed microfluidic device showed matching dimensions to the designed values.

### 2.2 Diameter-coded amphiphilic particles by tubing variations

The same 3D printed microfluidic device was connected to tubings of different *ID*_*t*_ to fabricate diameter-coded (*D*-coded) particles with clearly distinct outer diameters (*D*_*o*_) (**Figure 3**A). By using *ID*_*t*_ of 500μm (*D*500), 750μm (*D*750), and 1000μm (*D*1000), the *D*-coded amphiphilic particles with *D*_*o*_ of 334μm, 490μm, and 673μm and cavity diameters (*D*_*c*_) of 144μm, 239μm, 312μm were fabricated, respectively (Figure 3B). The slit length (*L*_s_ = 200μm) of 3D printed photomask (**Figure S2**) and the flow rate ratios (*O*1 = 1:1:1:1) remained constant in these experiments.

**Figure 3.**
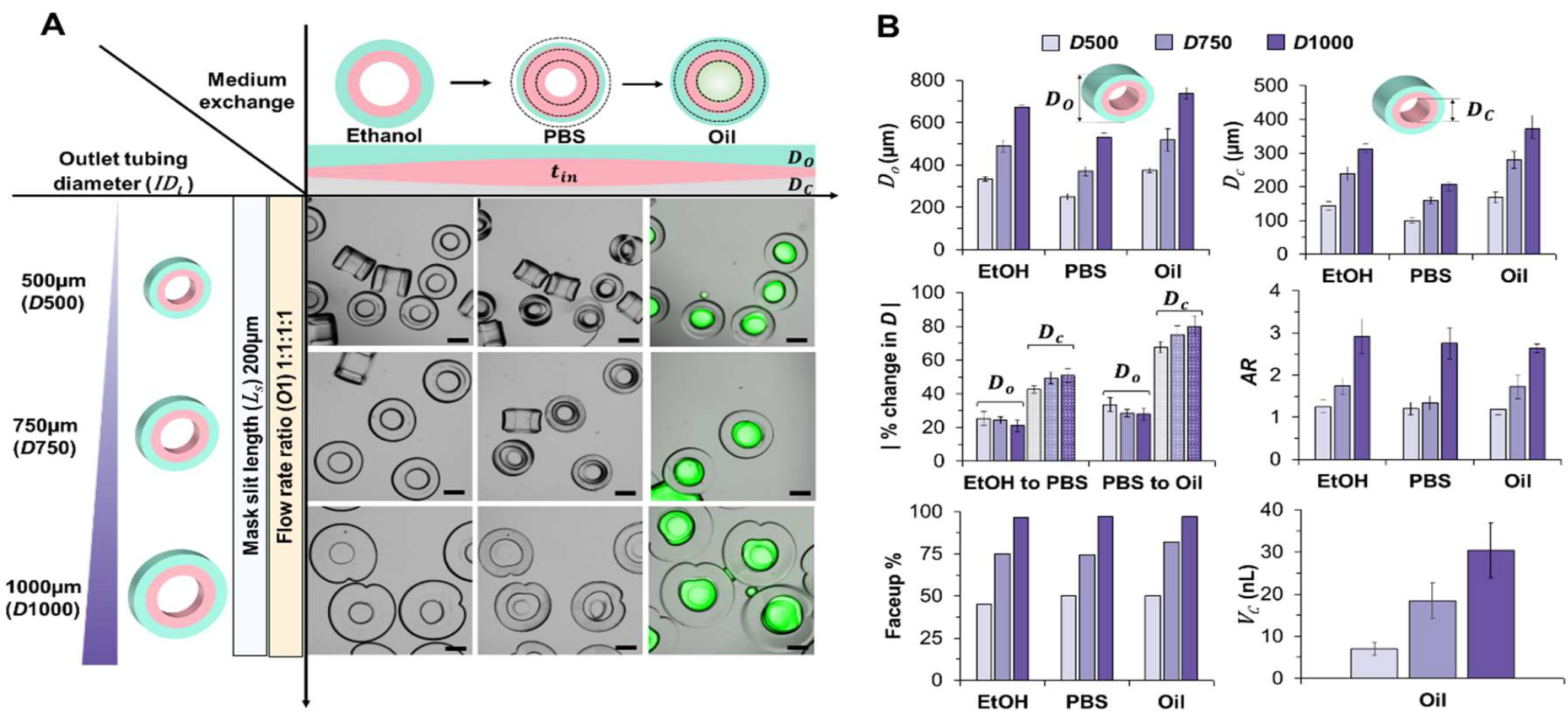
Diameter-based coding of concentric amphiphilic particles. (A) The fabrication of *D*-coded particles with *ID*_*t*_ of 500μm, 750μm, and 1000μm and with the same *L*_S_ (200μm) and *O*_1_ (1:1:1:1). The overall size (*D*_*o*_ and *D*_*c*_) of the particles decreased when the medium was exchanged from ethanol to PBS, and increased again when the medium was exchanged to oil (Scale bar 200μm). (B) Important parameters of *D*-coded particles (*D*500, *D7*50 and *D1*000) such as outer diameter (*D*_*o*_), cavity diameter (*D*_*c*_), percentage change in diameters (*D*_*o*_ and *D*_*c*_), aspect ratio (*AR*), faceup rate, and volume of cavity (*V*_*c*_), were measured in different media.

Overall, the particle dimensions (*D*_*o*_ and *D*_*c*_) increased with the increase in *ID*_*t*_, and a similar trend was observed with the exchange of medium from ethanol (EtOH) to PBS and oil. However, compared to a correlation coefficient of ∼0.66 between *D*_*o*_ and *ID*_*t*_ (i.e. *D*_*o*_ ≈ 0.66 × *ID*_*t*_) in ethanol, the average coefficient was recorded as ∼0.50 in PBS and ∼0.75 in oil. As the medium was initially exchanged from ethanol to PBS, the hydrophobic layer of the particles shrank resulting in a decrease of *D*_*o*_ by ∼25% and *D*_e_ by ∼35%. The hydrophobic layer swelled again when the medium was finally exchanged from PBS to oil, thereby, *D*_*o*_ and *D*_*c*_ increased by ∼30% and ∼45%, respectively.

The aspect ratio (*AR* = *D*_*o*_/*L*) of the particles influenced their orientation in the well plate. The fabricated *D*-coded particles had similar lengths (*L* ∼246μm) associated with slit length of *L*_s_ = 200μm, whereas the *AR* was calculated as ∼1.3, ∼2.0, and ∼2.7 for the *D*500, *D*750, and *D*1000 particles, respectively. As the *AR* increased, more particles faced upward in the well plate with the cross-sectional profile visible from the top. The faceup percentage increased from ∼40% for *D*500 (*AR* ∼1.3) particles to ∼80% and ∼95% for *D*750 (*AR* ∼2.0) and *D*1000μm (*AR* ∼2.7) particles, respectively. These results were consistent with the previous study showing more than 80% faceup rate for *AR*>2.5 for square shape particles ^16^.

The volume of a cylindrical particle cavity (*V*_*c*_) was calculated from measured particle dimensions (*D*_*o*_, *L*) as follows: *V*_*c*_ = *πD*_*c*_ ^2^*L*/4 . For the *D*-coded particles, the *V*_*c*_ was calculated to be ∼7nL, ∼19nL, and ∼30nL in oil for the *D*500, *D*750, and *D*1000 particles, respectively. The PBS droplet filling the cavity of the particle will have a volume equivalent to the *V*_*c*_.

### 2.3 Length-coded amphiphilic particles by mask variations

Modulating the masks with different slit lengths (*L*_S_), from 200μm to 2000μm (*L*2000), and with a minimum step size of 100μm, the following length-coded (*L*-coded) particles were fabricated: *L*200, *L*300, *L*500, *L*850, *L*1000, *L*1500, and *L*2000 (**Figure 4**A and **Figure S3**). The *L*-coded particles’ diameters (*D*_*o*_ and *D*_*c*_) showed a similar trend to that observed with *D-*coded particles suspended in different media.

**Figure 4.**
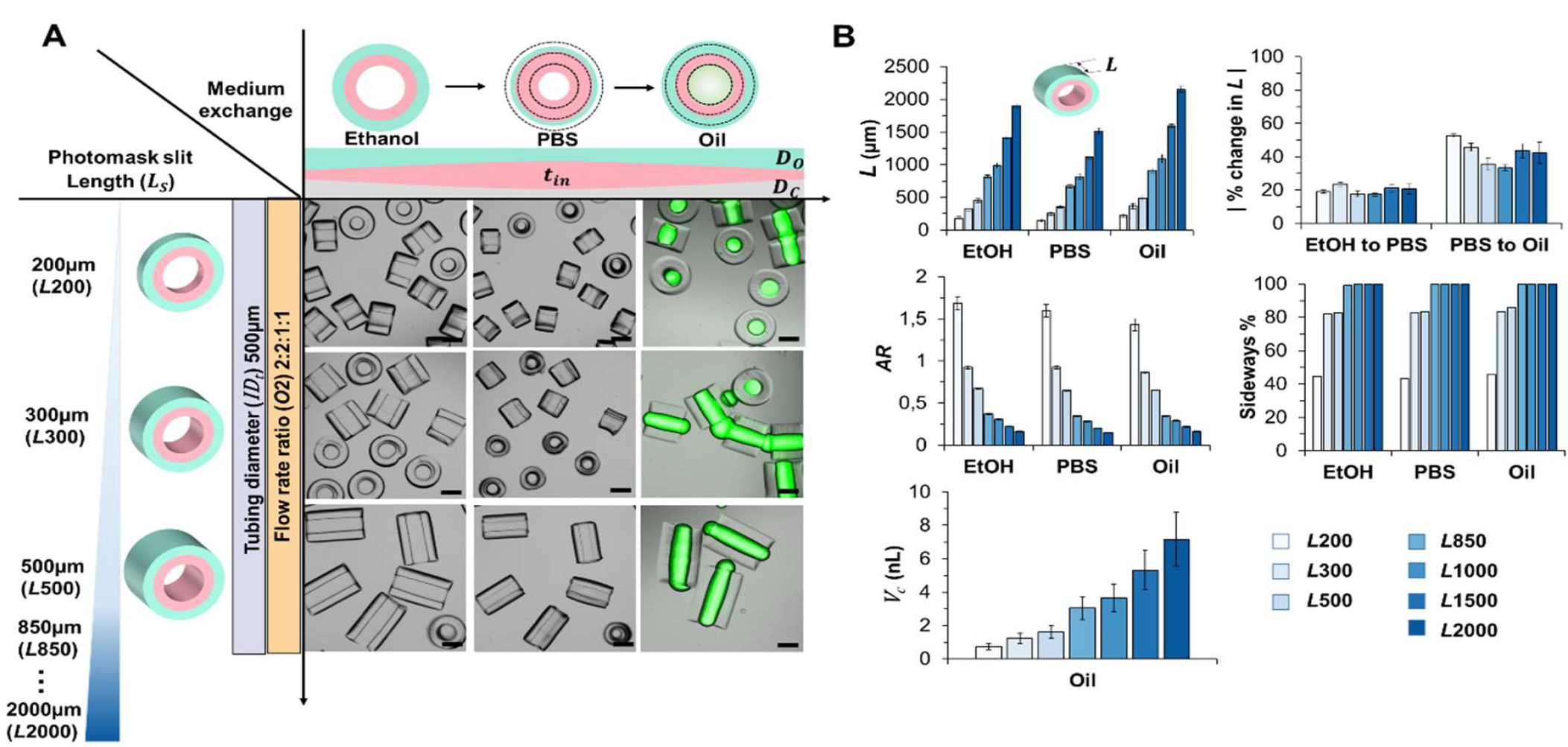
Length-based coding of concentric amphiphilic particles. (A) The *L*-coded particles were fabricated with the photomasks of varying *L*_S_ and with the same *ID*_*t*_ of 500μm and flow rate ratio of 2:2:1:1 (*O*2). The particle dimensions (*D*_*o*_, *D*_*c*_, and *L*) first decreased and later increased as the medium was exchanged from ethanol to PBS and from PBS to oil, respectively (Scale bar 200μm). (B) Important parameters, such as particle length (*L*), the percentage change in length (*L*) upon medium exchange, aspect ratio (*AR*), percentage of particles oriented in sideways directions, and the volume of cavity (*V*_*c*_) were measured and plotted for the *L*-coded particles.

Our stop-flow lithography setup included a UV light source composed of an array of multiple LEDs radiating a non-collimated light that necessitated 3D photomasks with up to 3cm height for segmented curing of precursor streams in the form of individual particles (Figure S2). The length of cured particles was within <15% of the slit length of the photomask (Figure 4B). The UV intensity and exposure time (*t*_*exp*_) were meticulously tuned to mitigate the negative effects of the overexposed precursor streams that would otherwise cure in the form of long fibers (**Table S2**). Moreover, the gap between the two adjacent slits of the 3D photomask was also adjusted to obtain distinct particles upon curing of discrete packets of the precursor streams. Larger slits were able to collect additional incident rays from oblique angles as the UV exposure area increased with the slit length; therefore, the exposure time and intensity were reduced for curing longer particles (see supporting information and Table S2).

The length of the *L*-coded particles on average decreased by ∼20% and subsequently increased by ∼25% as the medium exchanged from ethanol to PBS, and from PBS to oil, respectively. The *L*-coded particles, fabricated within an outlet tubing of *ID*_*t*_ = 500μm, had a constant diameter *D*_*o*_ of ∼300μm. The *AR* (= *D*_*o*_/*L*) of the particles decreased from ∼1.7 to ∼2.7 with an increase in the particles length from ∼178μm to ∼1902μm, which ultimately influenced the orientation of particles when seeded in the well plate. For the particles with lengths greater than 300μm (AR<1), more than 80% of the particles were facing sideways along the direction of their length. For *L* > 500μm, nearly all particles were seeded in the sideways orientation. The cavity volume *V*_*c*_ for *L*-coded particles fabricated with the *ID*_*t*_ of 500μm varied from ∼0.8nL for the smallest length particle (*L*200) up to ∼7nL for the longest particle length (*L*2000).

### 2.4 Engineering the amphiphilic particle size by flow rate ratio variations

#### 2.4.1 Tuning the cavity diameter

By modulating the flow-rate ratios of the polymer streams, the particles were fabricated with a range of inner cavity (*D*_*c*_) and outer particle (*D*_*o*_) diameters. In this experiment, the *L*_S_ (300μm), and *ID*_*t*_ (500μm) remained constant, while the flow rate ratios were varied in a systematic manner (**Figure 5**A). The flow rates of the outer two streams (*Q*_l,2_ = *Q*_1_ = *Q*_2_) and the inner two streams (*Q*_3,4_ = *Q*_3_ = *Q*_4_) were locked such that the ratio *Q*_l,2_: *Q*_3,4_ was varied as follows: 1:1(*O*1), 2:1(*O*2), 4:1(*O*4), and 8:1(*O*8). Similar *O*-coded particles were also fabricated by using the same microfluidic device connected with *ID*_*t*_ of 750 μm (**Figure S4**), and 1000 μm (**Figure S5**).

**Figure 5.**
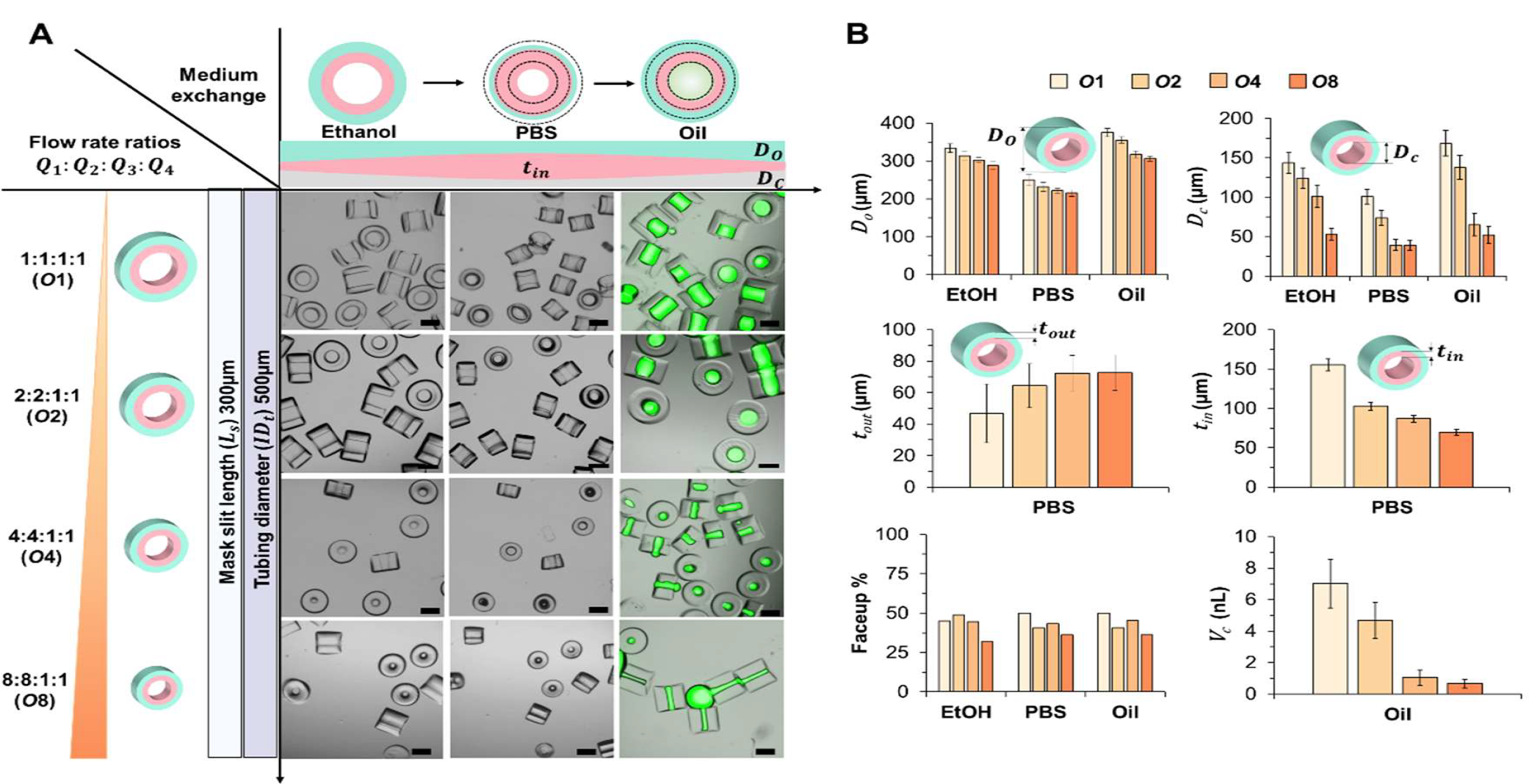
Controlling the cavity volume of concentric amphiphilic particles. (A) The *O*-coded particles were fabricated by varying flow rate ratios *Q*_1_: *Q*_2_: *Q*_3_: *Q*_4_, while *L*_s_ (300μm) and *ID*_*t*_ (500μm) remained constant. The flow rate ratio (*Q*_l,2_: *Q*_3,4_) increased from 1:1 to 8:1 to significantly reduce the cavity diameter of the fabricated particles from *O*1 to *O*8 (Scale bar 200μm). (B) Important parameters, such as outer diameter (*D*_*o*_), cavity diameter (*D*_*c*_), outer layer thickness (*t*_*out*_), inner layer thickness (*t*_*in*_), faceup rate, and cavity volume (*V*_*c*_), were measured and plotted for the *O*-coded particles.

As the flow rate ratio (*Q*_l,2_: *Q*_3,4_) was increased from 1:1 to 8:1, *D*_*o*_ (in ethanol) decreased by ∼13% (from ∼334μm to ∼288μm) while the *D*_*c*_ (in ethanol) decreased by ∼63% (from ∼154μm to ∼53μm) (Figure 5B). Numerical simulations of the cross-sectional flow profiles also confirmed a similar trend of particle dimensions (*D*_*o*_ and *D*_*c*_) with the flow rate ratio (Figure S1). The *D*_*o*_ of fabricated particles decreased by ∼20μm with each step increment in flow rate ratio from *O*1 to *O*8, which was relatively insignificant compared to the absolute change in *D*_*o*_ by >150μm from *D*500 to *D*750 for *D*-coded particles. Upon exchange of media from ethanol to PBS to oil, the *D*_*o*_ varied from ∼334μm to ∼249μm (-25%) to ∼375μm (+51%), and from ∼288μm to ∼215μm (-25%) to ∼305μm (+42%) for the *O*1, and *O*8 particles, respectively. Similarly, upon exchange of media from ethanol to PBS to oil, the *D*_*c*_ varied from ∼143μm to ∼101μm (-29%) to ∼167μm (+65%), and from ∼53μm to ∼39μm (-26%) to ∼52μm (+33%) for the *O*1, and *O*8 particles, respectively.

The thickness of the curable layers of the particles (*t*_*out*_, *t*_*in*_) was measured in the PBS as the layers were not distinct in the ethanol and oil phases. As the flow rate ratio (*Q*_l,2_: *Q*_3,4_) increased from *O*1 to *O*8, the thickness of outer particle layer (*t*_*out*_) increased (from ∼47μm to ∼73μm), whereas the thickness of inner particle layer (*t*_*in*_) decreased (from ∼155μm to ∼69μm).

With the increase in flow rate ratios from *O*1 to *O*8, the *AR* of the particle changed marginally from ∼1.1 to ∼0.96, thereby altering the faceup rate from ∼45% to ∼36%, respectively. The particle cavity volume (*V*_*c*_) decreased gradually (from ∼8nL to ∼0.8nL) with the increase in flow rate ratios as *D*_e_ of the particles decreased significantly by ∼63%. For the particles fabricated with the *ID*_*t*_ of 750μm and 1000μm, *V*_*c*_ decreased from ∼18nL (*O*1) to ∼1.6nL (*O*8) and ∼30nL (*O*1) to ∼2.4nL (*O*15), respectively (see Figures S4 and S5).

#### 2.4.2 Tuning the layer thickness

The flow rate ratio of the curable streams (*Q*_2_: *Q*_3_) was varied to modulate *t*_*in*_ and *t*_*out*_ as the *L*_S_ (300μm) and *ID*_*t*_ (500μm) remained constant (**Figure 6**A). The flow rate ratio of the inert streams was fixed (*Q*_1_: *Q*_4_ = 4:1), while the flow rate ratio of curable streams (*Q*_2_: *Q*_3_) was varied as follows: 4:2 (*O*4421), 2:2 (*O*4221), 2:4 (*O*4241), and 1:4 (*O*4141).

**Figure 6.**
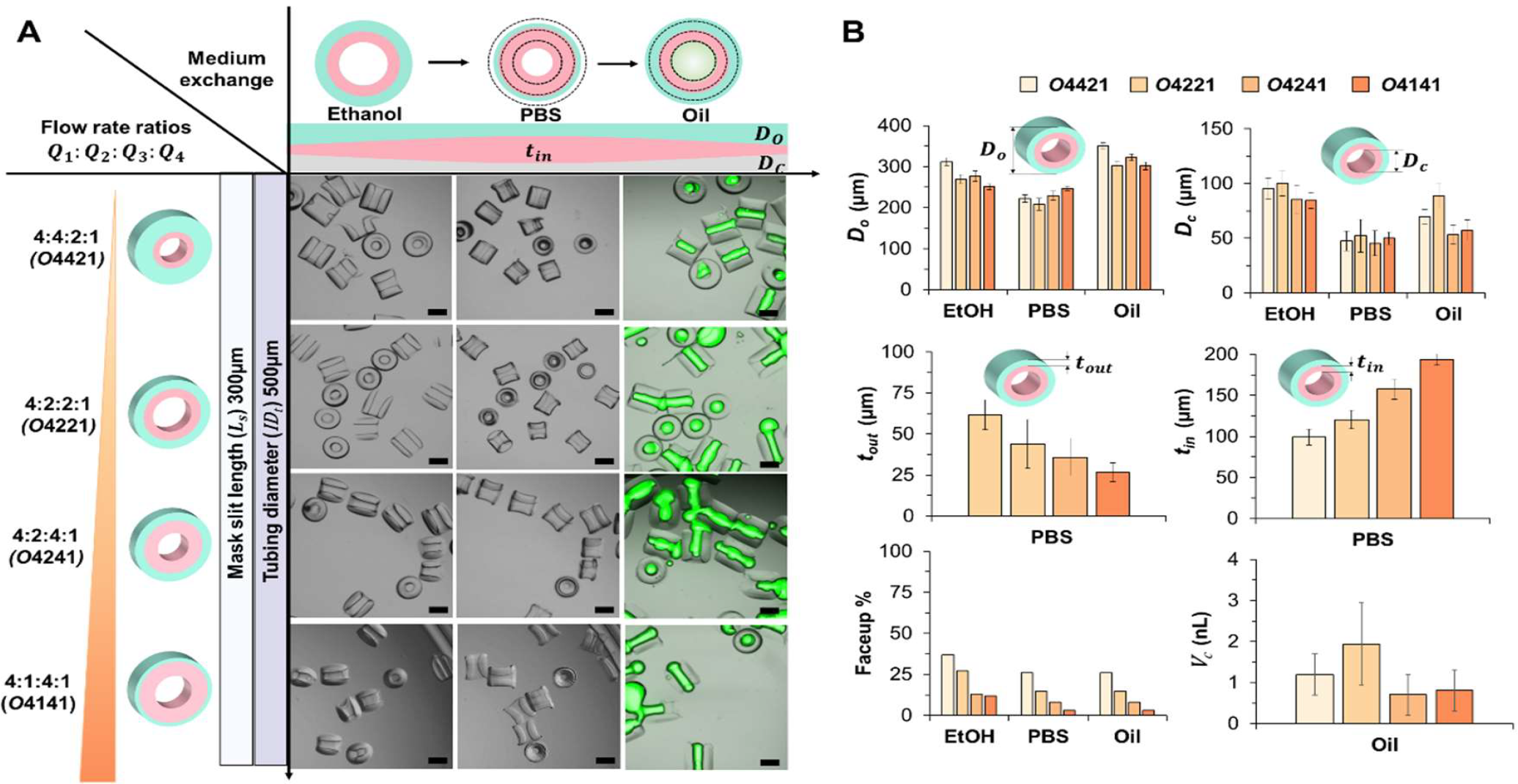
Controlling the layer thickness of concentric amphiphilic particles. (A) The O-shape particles were fabricated by varying flow rate ratios (*Q*_1_: *Q*_2_: *Q*_3_: *Q*_4_), while while *L*_s_ (300μm) and *ID*_*t*_ (500μm) remained constant. The flow rate ratio of the inert streams remained constant (*Q*_1_: *Q*_4_ =4:1), while the flow rate ratio of the curable streams (*Q*_2_: *Q*_3_) was tuned as 4:2, 2:2, 2:4, and 1:4 to obtain variable thickness of hydrophobic (*t*_*out*_) and hydrophilic (*t*_*in*_) layers of particle (Scale bar 200μm). (B) Important parameters, such as outer diameter (*D*_*o*_), cavity diameter (*D*_*c*_), outer layer thickness (*t*_*out*_), inner layer thickness (*t*_*in*_), faceup rate, and cavity volume (*V*_*c*_), were measured and plotted for the *O*-coded particles.

The *D*_*o*_ reduced by ∼24% from highest value of ∼311μm for *O*4421 to lowest value of ∼251μm for *O*4141, whereas the *D*_*c*_ decreased by ∼16% from highest value of ∼100μm for *O*4221 to lowest value of ∼84μm for *O*4141. The trends in *D*_*o*_ and *D*_*c*_ for different flow rate ratios was not consistent, as these dimensions were dependent on individual flow rates of the four precursor streams with non-uniform viscosities. The flow rates *Q*_1_and *Q*_2_ associated with higher viscosity outer precursor streams dominated the cross-sectional flow profile. As the flow rate ratio of the curable streams (*Q*_2_: *Q*_3_) was decreased from 4:2 (*O*4421) to 1:4 (*O*4141), *t*_*out*_ decreased by ∼57% (from ∼61μm to ∼26μm) and *t*_*in*_ increased by ∼94% (from ∼99μm to ∼193μm), which was consistent with our expectations.

The faceup rate of the particles decreased from ∼36% to ∼3% with the decrease in Do from ∼311μm (*O*4421) to ∼251μm (*O*4141). The cavity volume *V*_*c*_ ranged from ∼1.93nL (*O*4221) to ∼0.8nL (*O*4241) with the variation in flow rate ratios. With the increase in *t*_*in*_, more particles formed bridges due to stronger interaction of aqueous solution with the hydrophilic layer.

### 2.5 Size-based coding of particles

Amphiphilic particles were size-coded, i.e. *D*-coded and *L*-coded, by systematically configuring microfluidic device with different outlet tubings and photomasks, respectively (**Figure 7**A). Different dimensions of the size-coded particles were further engineered by tuning the flow rate ratios to vary the particle cavity volume. The size-coded particles were thoroughly characterized based on their dimensions, orientations, and abilities to hold uniform droplets within their cavities.

**Figure 7.**
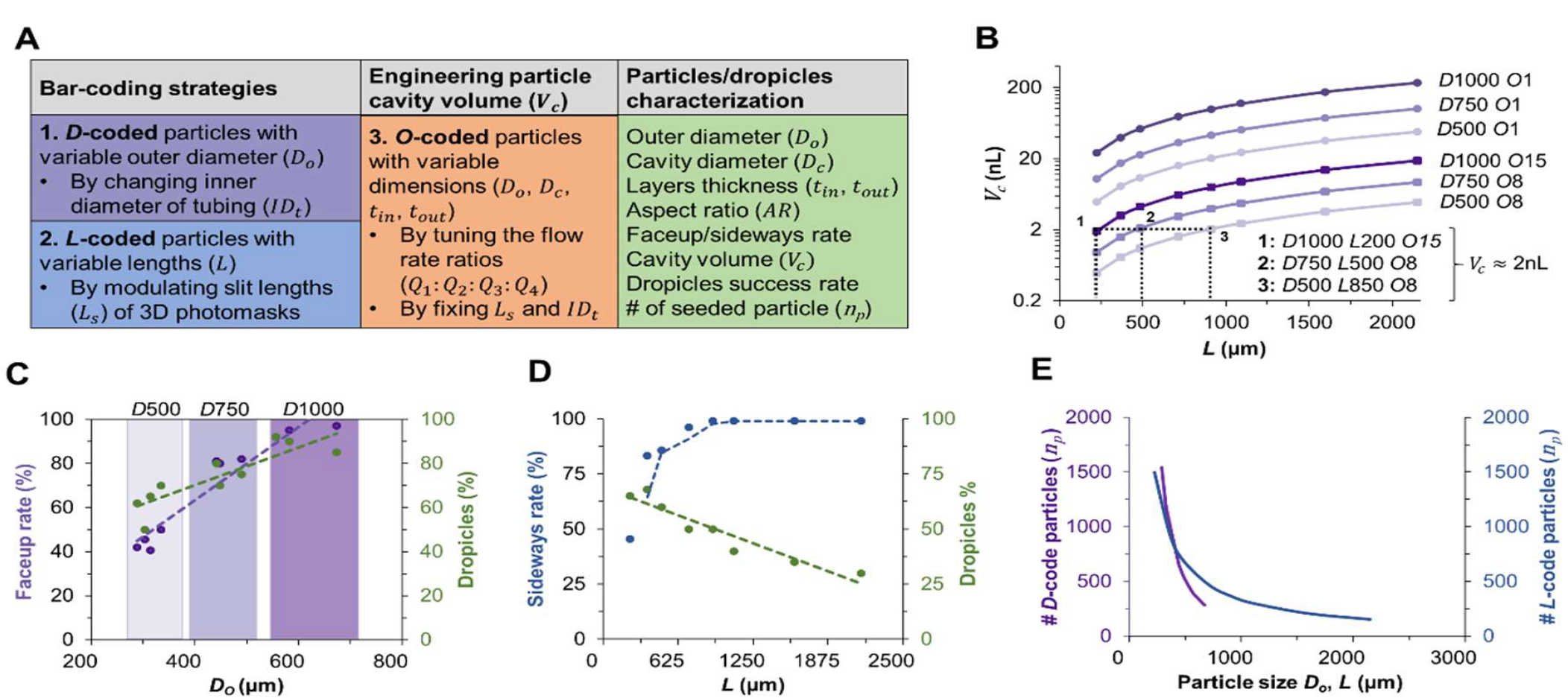
Relation between the size-coding strategy of amphiphilic particles and the particle/droplet characterization. (A) The size-coding strategy was mainly based on *D*_*o*_ and *L*, which were changed by varying the *ID*_*t*_ and *L*_s_, respectively. The flow rate ratios were adjusted to tune various particle dimensions (*D*_*o*_, *D*_*c*_, *t*_*in*_, *t*_*out*_) and the cavity volume (*V*_*c*_). The dropicles were characterized based on their dimensions, orientation, droplet formation, and seeding rate. (B) A range of *V*_*c*_ values were realized by combining various coding strategies. For a constant *V*_*c*_, it was possible to select unique size-coded particles. Orientation of the *D*-coded (C) and *L*-coded (D) particles is plotted together with the success rate of dropicle formation. (E) The number of particles (*n*_*p*_) that can be seeded inside a single well of the 24-well plate with 50% well coverage.

The cavity volume *V*_*c*_, estimated based on the cavity diameter and particle length, indicated that a wide range of droplet volumes (∼0.5-230nL) can be encapsulated within the amphiphilic particles (Figure 7B). In this work, we have demonstrated dropicle formation with volumes <30nL. It was observed that the longer particles (*L* >500) had difficulty filling up completely. For the same volume, multiple size-coded particles can be selected, such as ‘*D*1000-*L*200-*O*15’, ‘*D*750-*L*500-*O*8’ and ‘*D*500-*L*850-*O*8’ will have a similar volume of ∼2nL.

For the *D*-coded particles, the particle faceup orientation and the dropicle success rate increased linearly with the diameter. A dropicle filled up to 80% the cavity volume without forming aqueous bridges with neighboring particles was considered a successful count. The cavities of the smaller diameter particles were filling up with high percentage, however, due to lower faceup rate and aqueous bridges, the rate of segmented dropicles was lower. The particle openings facing each other prevented the droplets from splitting up in the absence of strong shear stress. The larger diameter particles could avoid the bridging problem due to higher face up rates, therefore, the dropicle success rate was >80% (Figure 7C). For the *L*-coded particles, the sideways orientation was >80% for *L* >300μm. The dropicle success rate dropped substantially to < 50% for particles with *L* >850μm as the trapped aqueous volume was not enough to fill the expanded cavity of the particles as the medium exchanged from PBS to oil.

We can seed up to 1500 particles with *D*_*o*_, *L* ≈ 300μm inside a single well of the 24-well plate with 50% area coverage. However, as the particle size increased for the *D*- and *L*-coded particles, the seeding density decreased sharply. For the *D*1000 and *L*2000 particles, the maximum seeding density was estimated as 288 and 154 particles per well, respectively.

## 3. Experimental

### 3.1 Design of 3D microfluidic device

The microfluidic device was designed by a computer aided design (CAD) software (PTC Creo Parametric, Student Edition) with an overall length of 27.5mm and width of 10mm (Figure 2). The device had four concentric channels to sculpt the flow in a circular cross-sectional profile. The inlet ports were designed with a tapered diameter of 1.65mm, which was reduced to 1.4mm for the tight insertion of polytetrafluoroethylene (PTFE) tubing (OD 1.59mm, ID 1.0mm, TECHLAB Germany) later on. The innermost channel of the device had a diameter of 0.8mm, while the other channels had a wall-wall spacing of 0.6mm and a wall thickness of 0.5mm in between the channels. The channels were supported by lateral pillar-like structures of 0.5mm diameter. The outlet port of the device had a diameter of 0.7mm. The device had a threaded outlet (1/4”-28 UNF) for connecting the transparent tubing using a flangeless fitting (IDEX flat bottom XP-235 PEEK Nut).

### 3.2 3D printing of microfluidic device and photomasks

The designed device was printed in-house by a stereolithography-based 3D printer (Phrozen Aqua Mini 8K, Phrozen Tech Co. Ltd., Taiwan) using a transparent resin (Anycubic Clear Resin). The CAD file was exported in stereolithographic (stl) format and was sliced (CHITUBOX Basic) in layers of 40μm thickness. The exposure parameters of 3D printing were tuned to obtain a fully enclosed 3D device without any blockage of channels. For the layer thickness of 40μm, the parameters were adjusted as follows: exposure time of 1.6s, bottom layer count of 4, and bottom layer exposure of 25s. A total of 6 devices were vertically printed on the build plate within a 3h cycle, consuming a resin volume of ∼15mL. The 3D photomasks were oriented along the build plate and printed with Aqua grey resin (Phrozen) with a 0.95s exposure time per layer with a thickness of 50μm. After 3D printing, the devices and photomasks were cleaned with isopropyl alcohol (VWR, Germany), and dried with nitrogen. After post-exposure in a UV bath for 45 minutes, four segments of PTFE tubing were inserted and glued into the four inlets of the 3D printed device.

### 3.3 Experimental setup for particle fabrication

The PTFE tubings (ID of 1mm, length of ∼55cm) connected the microfluidic device and the filled syringes (25mL glass syringes, Cetoni) by using flangeless fittings (XP-202 Delrin Nut, IDEX) and union connectors (P-603 standard unions, IDEX) on both ends. The syringes were mounted on four individually controllable syringe pumps (NemeSys S, Cetoni, Germany). The device was configured with a 10cm long transparent Teflon tubing (Perfluoroalkoxy or PFA) with the *OD*_*t*_ of 1.59mm and *ID*_*t*_ of 500μm, 750μm and 1000μm. A holder was printed (Ultimaker 3.0) to block the direct UV exposure of the microfluidic device, which also helped align the tubing in a straight line under the light source. A 3D printed photomask with an alignment holder was placed under the UV light source (Omnicure AC450, Excelitas Technologies, USA). Underneath of the 3D printed photomask, the outlet tubing with the structured fluid flow was selectively shined through the slits. The outlet tubing was connected with a pinch valve (2-way pinch valve, ASCO) on the other end through a soft tubing sleeve (ID1.57mm, OD3.18mm, hardness <55D). One end of the soft tubing sleeve was placed in the falcon tubing for particle collection. The syringe pumps, UV light source and the pinch valve were connected to an I/O-B module (Modular Qmix I/O-B Module, Cetoni) and were controlled by a custom script written in LabView software. The stop-flow lithography cycle (∼5s) was repeated multiple times to obtained ∼700 cured particles per minute in the collection tubes. The cured particles were washed with absolute ethanol for 2-3 times to completely remove the uncured precursors.

### 3.4 Stop-flow lithography for particle fabrication

The polymer precursors comprised of two inert streams (polypropylene glycol, PPG and polyethylene glycol, PEG) and two curable streams (polypropylene glycol diacrylate, PPGDA, and polyethylene glycol diacrylate, PEGDA), and were prepared according to the previously published protocols (details in the supporitng information). The precursor solutions were loaded in four individual syringes which were firmly held in the syringe pumps and connected to the microfluidic device (**Figure 8**A). The flow rate of each syringe pump (*Q*_1_ to *Q*_4_) was independently controlled by the LabView script. Polymer precursors with an inlety pressure of *P*_*in*_ at the syringes entered the 3D printed microfluidic device at pressure of *P*_d_. The device shaped the precuror streams and pushed them through the outlet transparent tubing at a pressure of *P*_*t*_ through a flangeless fitting. The structured streams were stopped momentarily as the pinch valve was closed and the syringe pumps were simulaneouly stopped. The outlet and inlet pressures were homogenized (*P*_*in*_ = *P*_*out*_) to remove the pressure gradient in the system, responsible for fluid flow. After a short delay time (*t*_d_), the streams were exposed to UV light by a rapidly switchable LED-based light source. The 3D-printed photomask collimated the incident light from the LED-array (Figure S2) to cure cylindarically-shaped amphiphlic particles. The cure particles were collected at the outlet by switching the pinch valve open and turning the syringe pumps on. A structured flow was developed again within stabalization time (*t*_s_) as the inlet pressure *P*_*in*_ reached a high value and the outlet pressure *P*_*out*_ dropped to the atmospheric level. The same cycle was repeated for multiple times continuous production of desired number of particles (Figure 8B). The pressure drop in the microfluidic device was obatained from the numerical simulation, where different inlets of the device were at different pressures due to the variable flow resistances of the individual channels (Figure S1B). However, the estimated pressure drop (flush phase) across the fluidic circuite, comprising of the inlet tubing (ID of 1mm), microfluidic device, and outlet tubing (*ID*_*t*_ of 0.5mm), indicated a minimal contribution by the microfluidic device (Figure 8C). Therefore, we can safely ignore the effect of mismached pressure drops in the device that could potential result in back flows within channels leading to undesirable mixing of the precursor streams during the stop phase.

**Figure 8:**
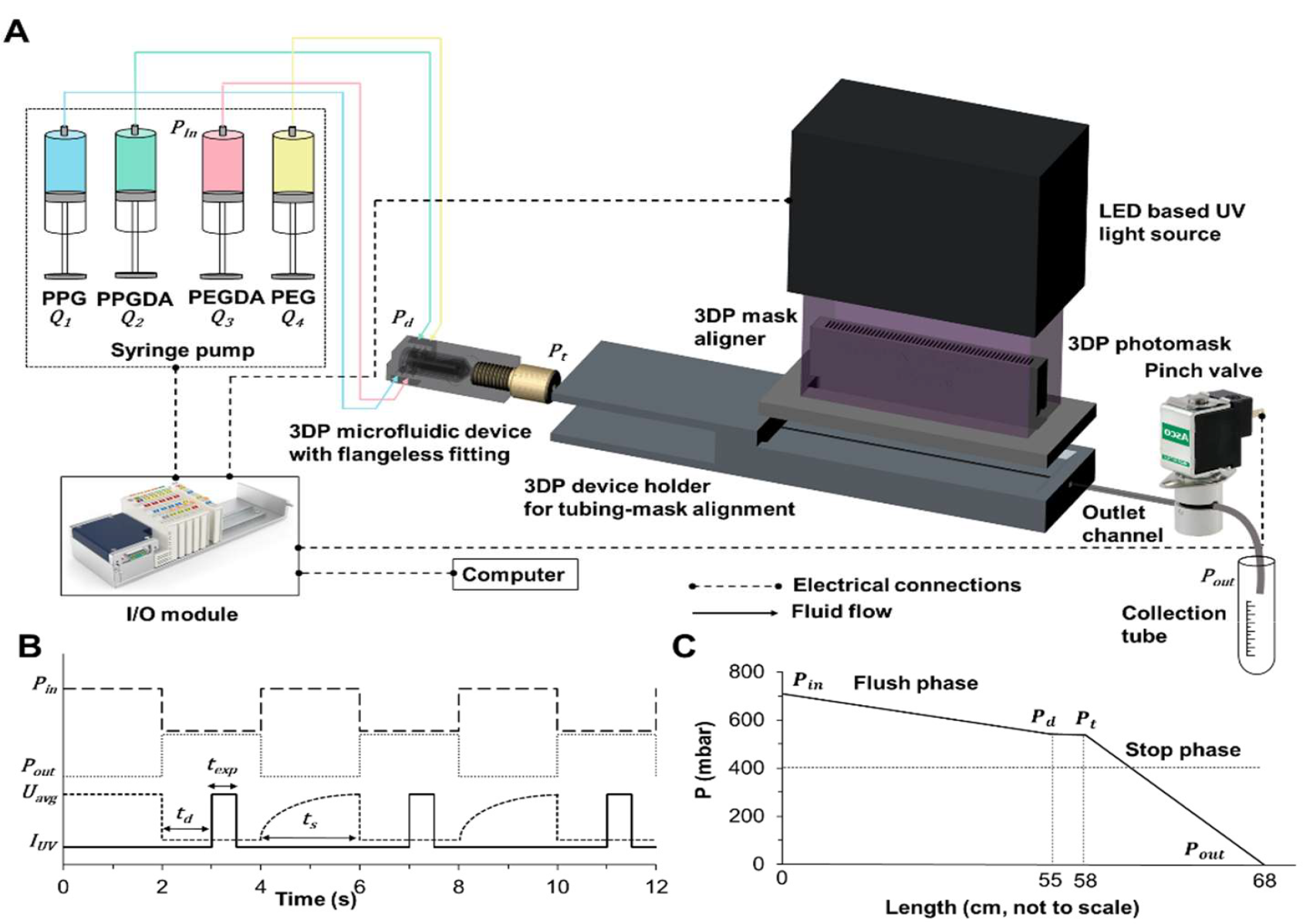
Stop-flow lithography setup for the fabrication of amphiphilic particles. (A) Syringes loaded with different precursors were mounted on the syringe pumps. Teflon tubings connected the syringes with the 3D-printed microfluidic device configured with a transparent tubing through the flangeless fitting at the outlet. A pinch valve was connected to the outlet tubing to regulate the fluid flow. A 3D printed holder aligned outlet tubing, microfluidic device, and 3D printed photomask under the UV light source. Pumps, light source, and pinch valve were controlled by a LabView script through an I/O module following a stop-flow cycle (B). (B) A pressure drop (*P*_*in*_ -*P*_*out*_) resulted in an average flow velocity of *U*_*avg*_. The flow was stopped by removing the pressure gradient to initiate the UV exposure for *t*_*exp*_ after a short delay *t*_*d*_. The flow was stabilized again within *t*_*s*_ as the pressure gradient was restored. (C) The pressure drops across the fluidic circuit with a minimal contribution by the microfluidic device.

### 3.5 Formation of dropicles

The fabricated particles were stored in ethanol after washing. The volume of ethanol was adjusted in a tube to obtain a specific concentration of the particles. The particles in ethanol were seeded in the hydrophobic well plate by using a pipette. The particles were settled at the bottom of the well due to a density difference, which allowed medium exchange without losing the particles after multiple washings. The medium was first exchanged from ethanol to aqueous solution (PBS), which was subsequently replaced with a fluorescent PBS solution for visualization purpose. As the excess aqueous solution was removed from the well plate, the droplets remained trapped inside the cavity of the particles. By the addition of oil (poly dimethylsiloxane-co-diphenylsiloxane or PSDS) in the well plate, segmented droplets within the particle cavities were formed.

## 4. Conclusion

We have demonstrated a configurable 3D printed microfluidics device for the fabrication of size-coded amphiphilic particles by varying the particle diameters (*D*-coded) or lengths (*L*-coded). A single device was reused multiple time with different outlet tubings and photomasks to produce a range of size-coded particles. Moreover, the flow rates of individual precursor streams were varied to readily tune the different dimensions of the particle, such as the outer diameter, cavity diameter, thicknesses of the hydrophilic and hydrophobic layers. The stop flow lithography setup incorporated an LED-based light source that was rapidly switched to cure particles without the need of a mechanical shutter. The 3D photomasks collimated the light originated from the LED-array, which was essential for the segmented curing of the particles. Finally, the amphiphilic particles, with a range of outer diameters (∼280μm to ∼670μm), lengths (∼180μm to ∼1900μm), cavity diameters (∼50μm to ∼310μm), hydrophilic layer (∼70μm to ∼190μm) and hydrophobic layer (∼26μm to ∼70μm) thicknesses in PBS, were fabricated. The fabricated particles could encapsulate droplets with ∼0.5nl to ∼30nL volumes in their cavities. The largest *D*-coded particles (*D*1000) had a faceup rate of >80% with a dropicle success rate of >95%. The *L*-coded particles mostly oriented sideways and had a dropicle success rate of <50%. However, the smallest L-coded particle (*L*200) had a sideways percentage of ∼40% (i.e. faceup rate of ∼60%) with a dropicle success rate of ∼65%. With a broad range of size-coded particles, we can encapsulate dropicles with similar volumes by modulating the diameter and length of the cavity.

## Supporting Information

### 1. Simulation of 3D microfluidic device

The laminar fluid flow inside the 3D printed devices was simulated by the “Laminar Flow” module in COMSOL Multiphysics for a stationary solution (Figure S1). The material properties of a single-phase were as follows: density 987 kg/m^3^, and viscosity 30mPa.s. The cumulative flow rate for the 4 inlets was fixed at 1.5mL/min, whereas the outlet boundary condition was set at the atmospheric pressure (0Pa). A concentric flow profile at the outlet of the tubing was obtained (Figure S1A). The outer diameter and the cavity diameter of the expected particle was reduced by increasing the flow rate ratios from 1:1:1:1 to 8:8:1:1. The pressure drop in the microfluidic device was measured for the four inlets. The inlets were at different pressures defined by the flow resistances of individual channels, with a maximum difference in pressure of ∼186Pa for given two inlets. The maximum pressure-drop (∼240Pa) across the microfluidic device was considerably low compared to the pressure drop across the inlet and outlet tubings (Figure S1B, and Figure 8C).

### 2. Materials preparation for particle fabrication

Four polymer precursors were prepared by mixing them in ethanol to match their densities by following our previous protocol. The outer inert hydrophobic stream, 90% polypropylene glycol (PPG, 1.01g/mL, M_w_ 400, Sigma-Aldrich) in 10% ethanol (0.79g/mL), provided a sheath flow to avoid clogging the tubing and defined the outer boundary of the particle. The inner inert hydrophilic stream, 60% polyethylene glycol (PEG, 1.12g/mL, M_w_ 200, Sigma-Aldrich) in 40% ethanol, defined the inner boundary of the particle in the form of a cavity. The curable hydrophobic stream, 90% polypropylene glycol diacrylate (PPGDA, 1.01g/mL, M_w_ 800, Sigma-Aldrich) in 10% ethanol, formed the outer layer of the particle upon UV exposure, while the curable hydrophilic stream, 60% polyethylene glycol diacrylate (PEGDA, 1.12g/mL, M_w_ 575, Sigma-Aldrich) in 40% ethanol, formed the inner layer of the particle. The prepared polymer solutions had a matched density of 0.987g/mL. A 5% (v/v) photoinitiator (2-hydroxy-2-methylpropiophenone, Darocur 1173, 405655, Sigma-Aldrich) was added in all four streams for photopolymerization of curable layers upon UV exposure.

### 3. Pattern transfer of the photomask upon UV exposure

Patterns from the 2D film and 3D printed photomasks were transformed to a curable PEGDA sample sandwitched between a glass slide and cover slip inside a 500μm deep well (Figure S2). The height of UV light source, fixed on a vertical translation stage (VAP 10, Thorlabs), was adjusted and the glass slide well was placed on the XY stage (XYT1/M, Thorlabs) to position the mask properly under the UV light source. The pattern was transferred from the 3D-printed photomasks to curable solution in the form of segmented slits. However, bulk of precursor was cured under the 2D mask without any impression of the slit pattern visible. By using variable slit gaps of 3D-printed photomasks, distinct patterns were formed. For the slit length of 200μm, and 10mm height of the photomask, the slit gap was varied as: 200μm, 500μm, and 1000μm. A 350μm wide pattern was obtained for a 200μm slit length and 500μm slit gap, which showed that the extra cured region (∼75μm on each side) was because of the non-colimated light travelling obliquely to the curable precursor. The height of photomask was increased to 15mm for 200μm slit length to obtain the pattern within ±20μm range of the designed slit length. As the slit length increased in the photmask for length-coded particles, the height was increased to 20mm for 300μm slit length and 30mm for 500μm slit length to obtain particles within ±40μm range (see Table S2 for additional detail).

### 4. Experimental parameters for particle fabrication

The polymer streams were pumped individually with a specific flow rate (*Q*_1_, *Q*_2_, *Q*_3_, *Q*_4_), but they were adjusted in such a way that the overall flow rate of the four streams remained close of 1.5mL/min. Different outlet tubings were configured with the device (500μm, 750μm, and 1000μm) while the overall flow rate remained the same. The UV light intensity (*I*_UV_) was ∼3.36W/cm^2^ at 4V (input voltage). The exposure time (*t*_*exp*_) of 1s was used for a photomask with minimum length of 200μm. The UV intensity remained similar, but the exposure time (*t*_*exp*_) was decreased with the increase in the length of the photomask. For the 500μm slit length of photomask, *I*_UV_ of 4V and *t*_*exp*_ of 500ms were used. The delay time (*t*_d_) during the stop-flow cycle was also controlled after applying pressure at the outlet through pinch valve to ensure that the flow was fully stopped before UV exposure. The delay time (*t*_d_) for the outlet tubing of 500μm, 750μm, and 1000μm was adjusted to 1s, 800ms and 500ms, respectively. The corresponding stabilization times (*t*_s_) were 3s, 4s, and 5s for the three tubing diameters, 500μm, 750μm, and 1000μm, respectively.

### 5. Dropicles imaging and characterization

A specific volume of the fabricated particles suspended in ethanol (usually 350μL for the particles with concentration of ∼700particles/ml) were seeded in the center of hydrophobic well plate (Falcon 24-well plate, 351147 Corning) by using a 1mL pipette. For loading the bigger particles (dimension >600 μm) in the well plate, the pipette tip (1mL) was cut to enlarge the opening. After seeding the particles (in ethanol) in individual wells, a bright field image of each well was captured under the microscope (Leica DMi8 inverted microscope). By exchanging the medium from the ethanol to PBS, the image was captured again. After the encapsulation of flourescent droplet in the cavity of particles, an overlay image of the particle with flourescent and brightfield channels was captured. The particles were analyzed in ImageJ manually to characterize the dimensional parameters.

**Table S1.** Cost estimation for a 3D printing of microfluidic devices and photomasks.

| 3D Printing | Material | Material volume<br>(each device per<br>print) | Material cost<br>(per device) | Cleaning cost with<br>IPA (per device) |
| --- | --- | --- | --- | --- |
| Microfluidic<br>device | Anycubic Aqua<br>clear | 4mL | 0.2\$ | 0.5\$ |
| Photomask | Phrozen Aqua<br>Gray | 5mL | 0.45\$ | 0.5\$ |

**Table S2.** Important experimental parameters.

| # | Size-<br>code | $ID_t$<br>($\mu\text{m}$) | Photomasks dimensional<br>parameters | | | Flow<br>rate<br>ratios | Total<br>flow<br>rate<br>($\text{ml}/\text{min}$) | Stop-flow Lithography Parameters | | | |
| --- | --- | --- | --- | --- | --- | --- | --- | --- | --- | --- | --- |
| | | | Slit<br>Length<br>$L_s$ ($\mu\text{m}$) | Slit<br>gap<br>$G$<br>($\mu\text{m}$) | Height<br>$H$<br>(cm) | | | Delay<br>time<br>($t_d$)<br>(ms) | Stabalization<br>time ($t_s$) (ms) | Expos-<br>ure<br>time<br>($t_{\text{exp}}$)<br>(ms) | Voltage<br>(V)<br>~ UV<br>intensity<br>(%)* |
| 1 | D500 | 500 | 200 | 500 | 1.5 | O1, O2,<br>O4, O8 | 1.35-1.5 | 1000 | 3000 | 1000 | 4 (80%) |
| 2 | D750 | 750 | 200 | 500 | 1.5 | O1, O4,<br>O8 | 1.35-1.5 | 800 | 4000 | 1000 | 4 (80%) |
| 3 | D1000 | 1000 | 200 | 500 | 1.5 | O1, O4,<br>O15 | 1.35-1.8 | 500 | 5000 | 1000 | 5 (100%) |
| 4 | L200 | 500 | 200 | 500 | 1.5 | O2 | 1.5 | 1000 | 3000 | 1000 | 4 (80%) |
| 5 | L300 | 500 | 300 | 500 | 2.0 | O2 | 1.5 | 1000 | 3000 | 850 | 4 (80%) |
| 6 | L500 | 500 | 500 | 500 | 3.0 | O2 | 1.5 | 1000 | 3000 | 500 | 4 (80%) |
| 7 | L850 | 500 | 850 | 500 | 3.5 | O4 | 1.5 | 1000 | 3000 | 500 | 3 (60%) |
| 8 | L1000 | 500 | 1000 | 500 | 4.0 | O4 | 1.5 | 1000 | 3000 | 500 | 3 (60%) |
| 9 | L1500 | 500 | 1500 | 500 | 4.5 | O4 | 1.5 | 1000 | 3000 | 500 | 3 (60%) |
| 10 | L2000 | 500 | 2000 | 500 | 5.0 | O4 | 1.5 | 1000 | 3000 | 500 | 3 (60%) |

\*Maximum rated UV intensity of the light source provided by the manufacturer was $8\text{W}/\text{cm}^2$ .

**Figure S1.**
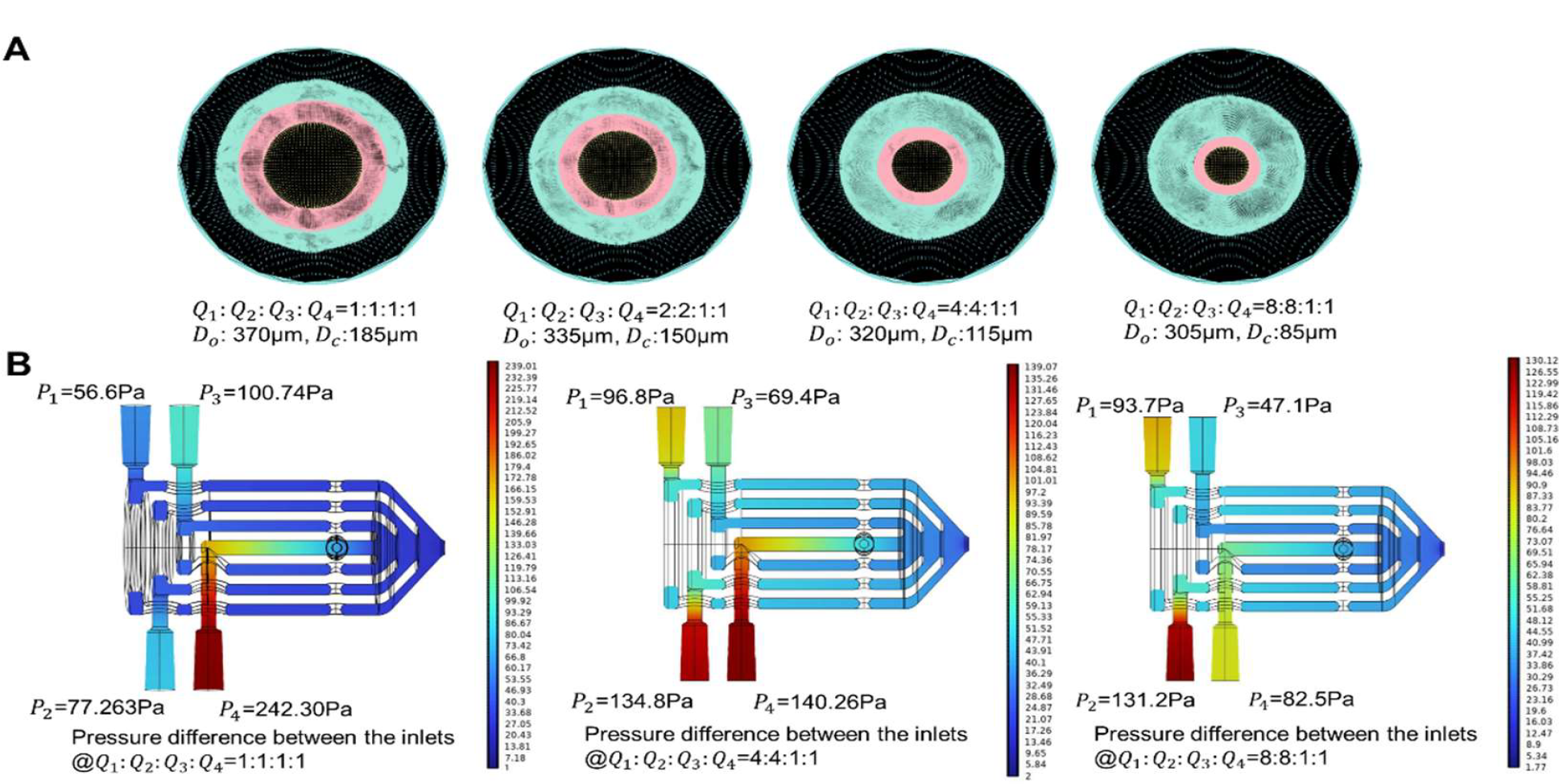
Laminar single-phase COMSOL simulation of the microfluidic device. (A) Fluid flow profile in the microfluidic device configured with tubing of 500μm diameter with increasing flow rate ratios from 1:1:1:1 to 8:8:1:1. (B) Pressure drop in the center plane of microfluidic device at increasing flow rate ratios.

**Figure S2.**
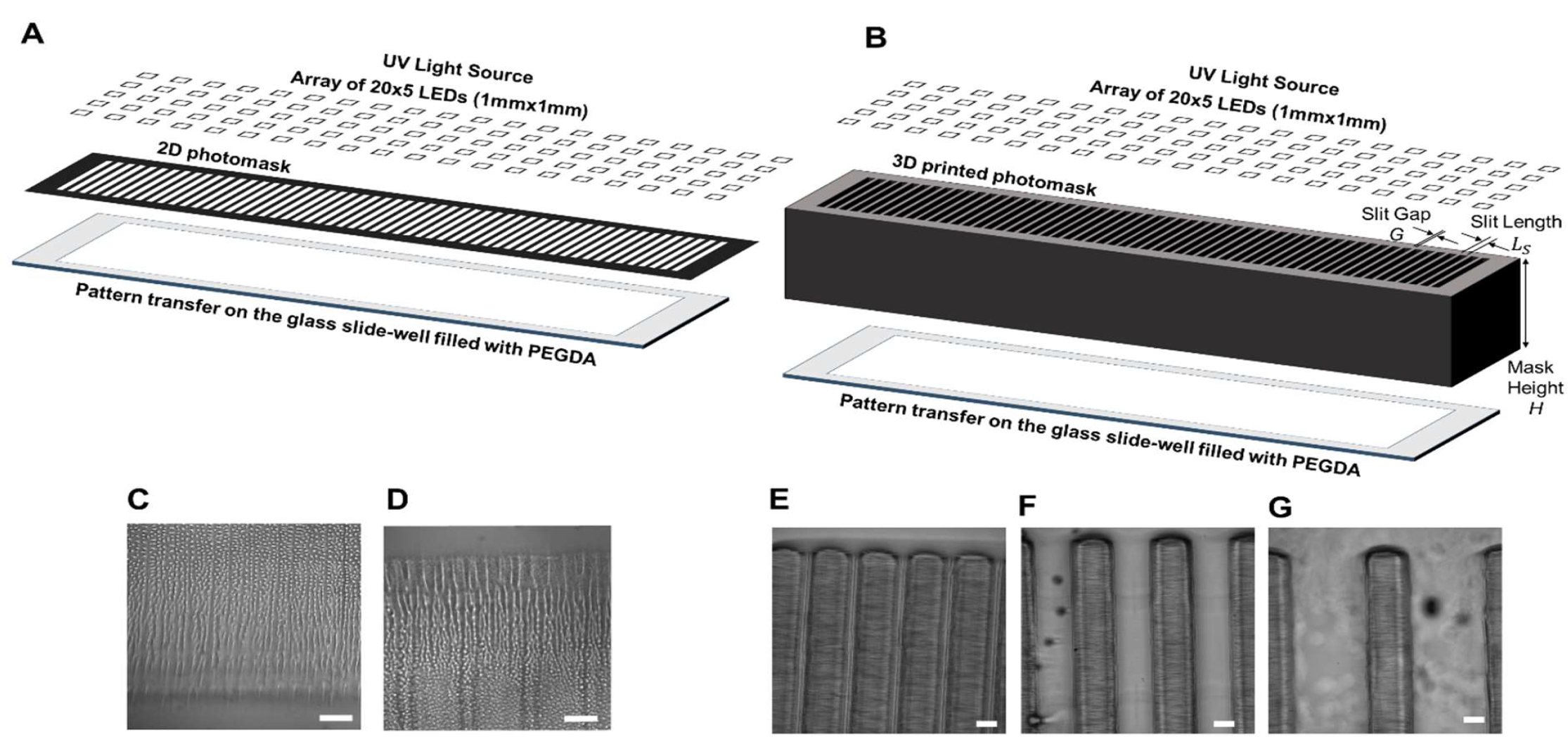
UV curing of PEGDA in a sandwiched well using a 2D and 3D-printed photomasks. (A, B) Schematics of pattern transfer of the slits present on the 2D and 3D photomask onto the glass slide filled with PEGDA after exposing UV light from the LED based light source. The photomask was placed directly on top of the glass slide while the UV light source had a gap of 20cm from the photomask. (C, D) Overexposure under the 2D photomask resulted in indistinguishable pattern due to non-collimated light from the source (Scale bar 200μm). (E, F, G) Pattern transfer from a 3D photomask with height of 10mm and slit length of 200um, and varying slit gap to block the light coming at an angle (Scale bar 200μm). (E) Slit gap 200μm. (F) Slit gap 500μm. (G) Slit gap 1mm.

**Figure S3.**
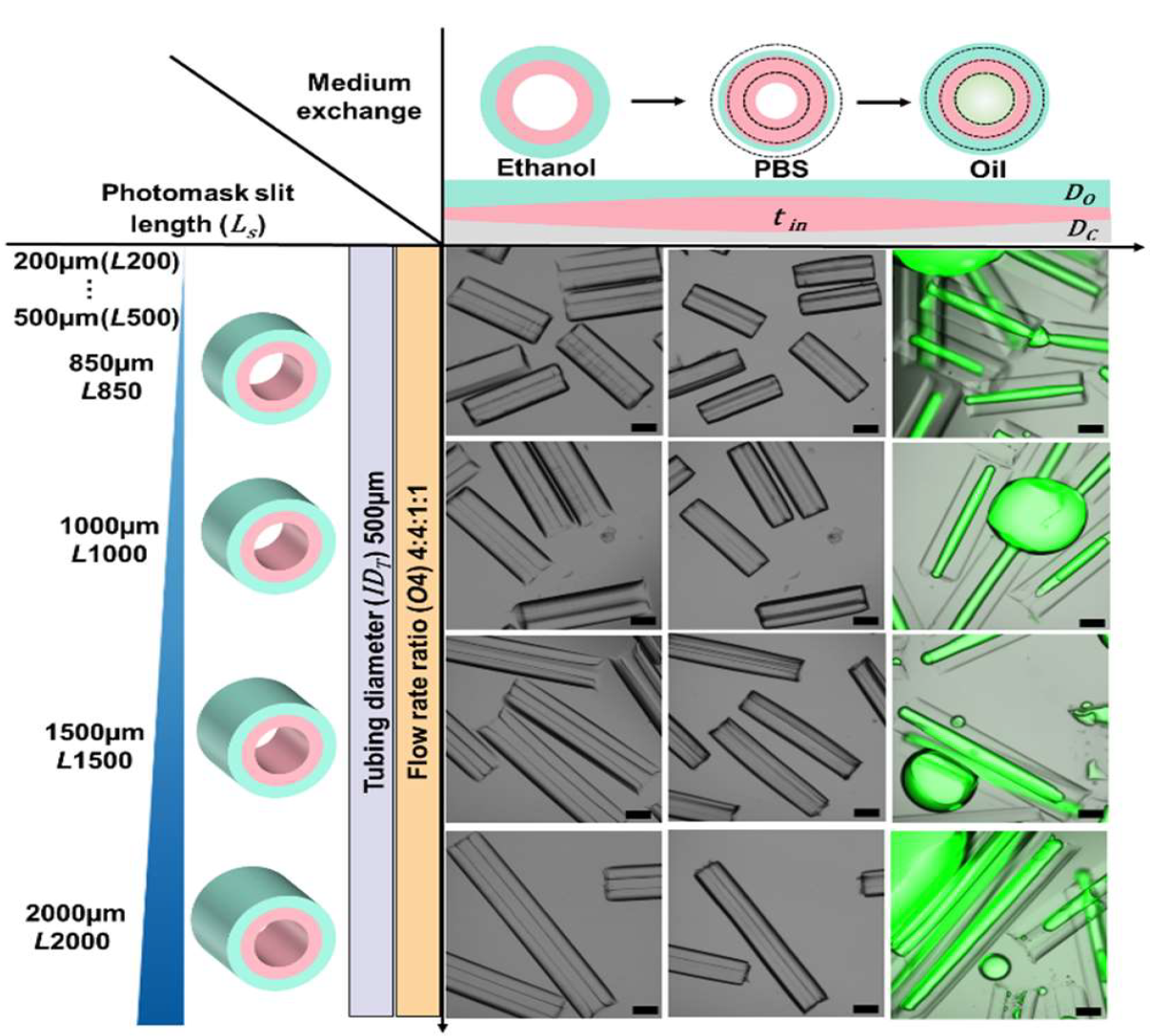
*L*-coded amphiphilic particles (850μm-2000μm). The fabrication of length-coded particles with the photomasks of varying *L*_s_ and with the same *ID*_*t*_ of 500μm and same flow rate ratios of *O*4.

**Figure S4.**
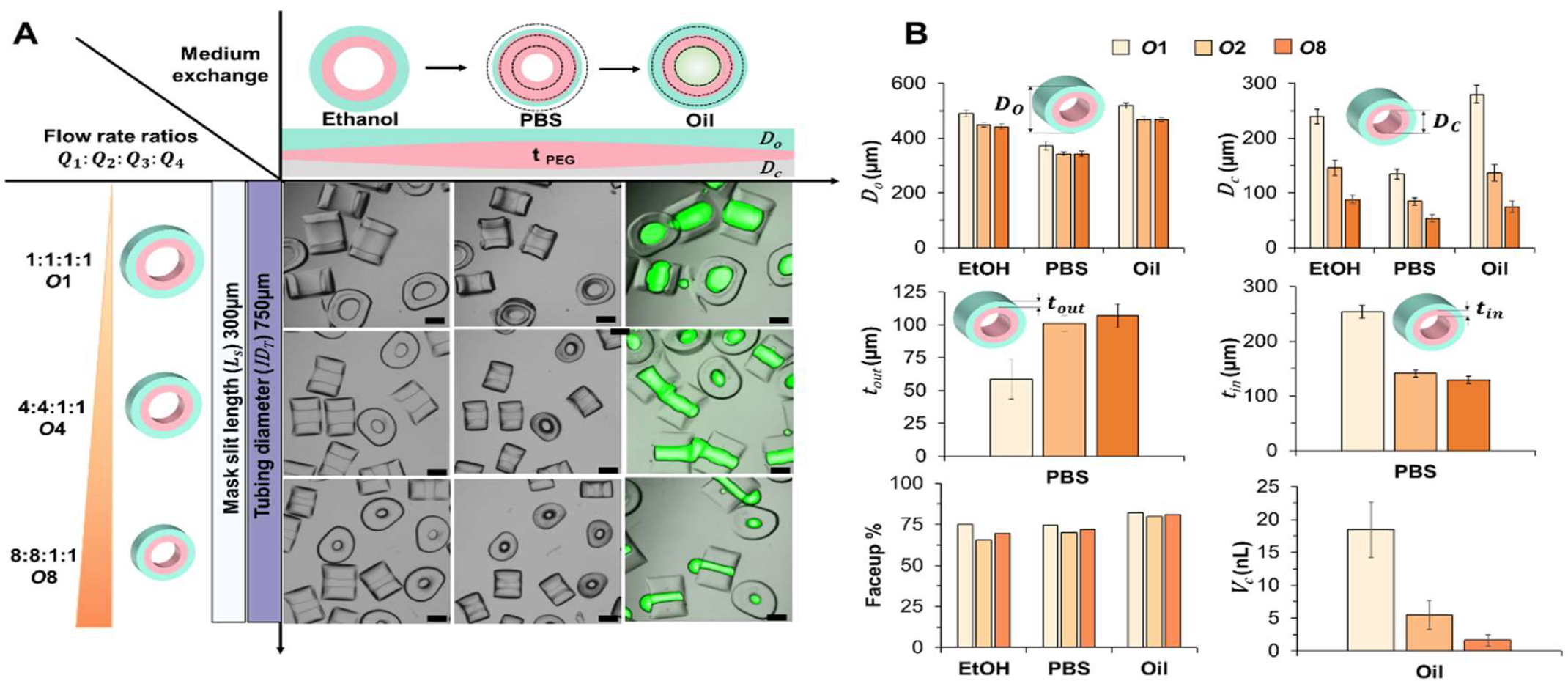
Controlling the cavity volume of concentric amphiphilic particles fabricated within the *ID*_*t*_ of 750 μm. (A) The fabrication of particles with varying flow rate ratios (*Q*_1_: *Q*_2_: *Q*_3_: *Q*_4_), where *Q*_1_= *Q*_2_ and *Q*_3_= *Q*_4_, while the *L*_*s*_ = 300μm and *ID*_*t*_ = 750μm. The flow rate ratios increased from 1:1 to 8:1 to reduce the cavity size of particles. (B) Particle cross-sectional dimensions such as outer diameter (*D*_*o*_), cavity diameter (*D*_*c*_), PPG layer thickness (*t*_*out*_), PEG layer thickness (*t*_*in*_), faceup rate, and the volume of cavity (*V*_*c*_) were plotted.

**Figure S5.**
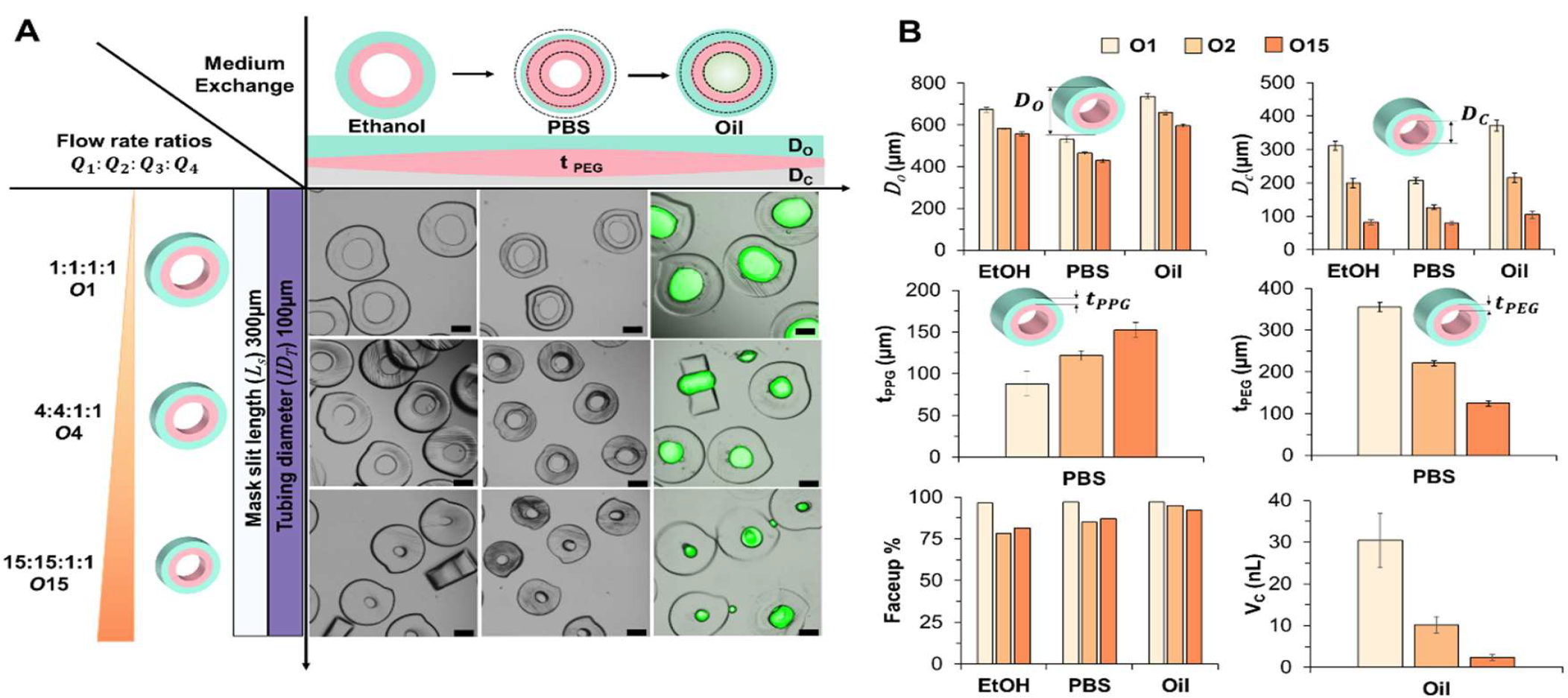
Controlling the cavity volume of concentric amphiphilic particles fabricated within the *ID*_*t*_ = 1000 μm. (A) The fabrication of particles with varying flow rate ratios (*Q*_1_: *Q*_2_: *Q*_3_: *Q*_4_), where *Q*_1_= *Q*_2_ and *Q*_3_= *Q*_4_, while the *L*_*s*_ = 300μm and *ID*_*t*_ = 750μm. The flow rate ratios increased from 1:1 to 15:1 to reduce the cavity size of particles. (B) Particle cross-sectional dimensions such as outer diameter (*D*_*o*_), cavity diameter (*D*_*c*_), PPG layer thickness (*t*_*out*_), PEG layer thickness (*t*_*in*_), faceup rate, and the volume of cavity (*V*_*c*_) were plotted.

